# The histone code of love: epigenetics of maturation of gonads in the human blood fluke *Schistosoma mansoni*

**DOI:** 10.64898/2026.01.23.701327

**Authors:** Christoph Grunau, Zhigang Lu, Avril Coghlan, Max Moescheid, Thomas Quack, Cristian Chaparro, Eerik Aunin, Jean-Francois Allienne, Adam Reid, Nancy Holroyd, Matt Berriman, Gilda Padalino, Karl F. Hoffmann, Christoph G. Grevelding, Ronaldo de Carvalho Augusto

## Abstract

*Schistosoma mansoni* is a parasitic flatworm that has two, genetically determined, sexes. We used aggregated data of 8 posttranslational histone modifications (ChIP-Seq), chromatin accessibility (ATAC-Seq), transcription (RNA-Seq) and genome feature annotations to decipher the histone code of genes involved in the differentiation of schistosome gonads (i.e. female ovaries and male testes). We show schistosome gonads express at least two classes of protein coding genes: H3K4me3-positive genes that display canonical features of eukaryotic protein-coding genes such as peaks of H3K4me3 at the transcription start sites (TSS) and increases in histone acetylation marks towards the transcription end site (TES), but also a non-canonical H3K9/27me3 plateau just upstream of the TSS. H3K4me3 enrichment at the TSS is highly predictive for transcription strength in these genes compared to a second class of protein coding genes (H3K4me3-negative genes) that do not display this pattern and is characterised by absence of the investigated histone marks at TSS and TES. This is indicative of the existence of hitherto unknown, potentially schistosome-specific histone marks in these genes. The absence of H3K4me3 at the TSS is not associated with inducible or stable gene expression in the gonads. Instead, gene ontology analysis indicates that H3K4me3-positive genes are related to functions which typically govern processes such as metabolism or signal transduction while H3K4me3-negative genes are dedicated to cell communication or immune responses. Second, individual histone modifications and their combinations are associated with functional features of the schistosome genome, known as “chromatin colours”. In H3K4me3-positive genes, there is clear co-linearity of 3 colours, which strongly suggests a functional role for histone modifications in the control of transcription pre-initiation, promotor release, and transcription termination. Third, there are striking chromatin structure changes during maturation of the gonads in all genomic features including protein-coding and non-protein coding genes as well as repetitive sequences. The nature of these changes is different in both sexes. H3K36me3 and H3K9me3, as well as H3K23ac and H3K9ac show the strongest variations. Last, we show that pharmacological inhibition of histone demethylation activity by IOX1 leads to a concentration-dependent separation (“divorce”) of schistosome couples confirming the importance of H3K36/H3K9 methylation for pairing maintenance and indicating histone demethylases as a potential drug target family. Collectively, our findings offer unprecedented insights into histone codes and chromatin dynamics governing the reproductive development of *S. mansoni* gonads.

## Introduction

Schistosomiasis is one of the most severe parasitic diseases affecting humans with at least 240 million requiring preventative treatment (Mutapi et al., 2017). Schistosomes have a complex life cycle with freshwater snails as the intermediate hosts and humans among the definitive hosts. However, several mammals can act as reservoirs in natural epidemiological sites and, thus, have an additional role in the development of control strategies (Huang and Manderson, 2005). Once inside the mammalian host, schistosomes migrate to their final destinations in the mesenteric veins or the urinary venous plexus where they start daily production of 300–3,000 eggs per female, depending on the species (Cheever et al., 1994). The pathology of schistosomiasis is triggered by eggs that do not reach the gut lumen (for *S. mansoni* and many other species) or the bladder (for *S. haematobium*), representing approximately half of the total egg production. These eggs eventually lodge in organs such as liver, where they penetrate the tissues causing severe inflammation and liver cirrhosis, the main cause of mortality (Colley et al., 2014; Olveda et al., 2014; McManus et al., 2018). Beyond its medical importance, the reproductive biology of schistosomes is an exceptional model to study sexual polymorphisms: among digenetic flukes, schistosomes possess the sex chromosomes ZW, which are not found in the other groups of parasitic flukes that are hermaphrodites. In addition, schistosomes show strong developmental polymorphism with striking morphological and physiological changes. After pairing, tri-directional molecular communication between male and female worms and the host (Moescheid et al., 2025) is require to produce mature egg-producing females. Female sexual maturation depends on constant pairing contact with the male, which induces substantial gonad-associated genes changes (Lu et al., 2016) as a prerequisite to achieve gonad differentiation together with host factors. Historically, the development of male schistosomes was considered independent of pairing since they possess fully mature testes and functional seminal vesicles even before finding their partners (Fitzpatrick and Hoffmann, 2006; Beckmann et al., 2010; Brandão-Bezerra et al., 2019). Recent data have shown an effect of pairing also on male’s gonad-associated genes, even though no strong morphological differences between male with and without pairing experience exist (Neves et al., 2005; Beckmann et al., 2010; Lu et al., 2016; Lu et al., 2019).

Molecular mechanisms underlying the mutual influence were only recently discovered and support our earlier hypothesis (Lu et al., 2019) that distinct, parallel regulatory systems coordinate female sexual maturation. For example, physical contact with the female induces expression of *gli1* mRNA in male which in turn upregulates *S. mansoni* nonribosomal peptide synthetase (*Sm-nrps*, Smp_158480) expression in the ciliated sensory neurons, located throughout the gynecophoral canal. *Sm-*nrps catalyzes selective coupling of the decarboxylation product of tryptophan, tryptamine, and ß-alanine to yield the ß-alanyl-tryptamine dipeptide (BATT) (Chen et al., 2022). BATT is secreted from paired male schistosomes throughout their gynecophoral canal and acts as a tigger for sexual development on the female. However, this is not the only signal: RNAi experiments have revealed essential roles for *Smtdc-1* (L-tyrosine decarboxylase, Smp_135230) and *Smddc-1* (aromatic-L-amino-acid decarboxylase, Smp_171580). Both genes are expressed in a pairing-dependent and male-specific (*Smtdc-1*) or male-preferential (*Smddc-1*) manner. Their knockdown led to pronounced phenotypic effects on the gonads and egg production in females, including impaired differentiation of mature oocytes (Li et al., 2023). Given that these genes are strictly regulated by pairing status—being transcriptionally activated in a characteristic “on-off-on” pattern during pairing, separation, and re-pairing—they may serve as male-derived competence factors.

After pairing, transcriptional changes of around 3,600 genes occur in the ovaries, and the contact with the female has also a profound effect on expression of 243 genes in the male gonads (Lu et al., 2016).

In schistosomes, each developmental stage is characterised by specific patterns of posttranslational histone modifications, which are also involved in an adaptive response (Roquis et al., 2018; Augusto et al., 2019; Picard et al., 2019; Fneich et al., 2016; Picard et al., 2016). We reasoned that histone modification might also be involved in sexual maturation and decided to analyse this in *S. mansoni* gonads. The local composition of chromatin is a major determinant of the transcriptional activity of a gene, and it was first proposed in fruit flies that post-translational modifications (PTM) of histones are part of a “chromatin colour-code” that “paint” the genome to indicate different activity levels (Filion et al., 2010). To decipher the chromatin colour-code of *S. mansoni* gonads, we choose a broad panel of eight key histone modifications with a theoretical yield of 256 colours (combinations of histone marks): H3K4me3, H3K9me3, H3K9ac, H3K23ac, H3K27me3, H3K36me3, H4K12ac, and H4K20me1. These histone PTMs were selected since they had been reported to influence developmental plasticity in other animal models. The roles of tri-methylated lysine 4 of histone H3 (H3K4me3), and acetylated lysine 9 of histone H3 (H3K9ac) are generally associated with euchromatin DNA regions (relaxed DNA, allowing transcription). Tri-methylated lysine 27 of histone H3 (H3K27me3), and tri-methylated lysine 9 of histone H3 (H3K9me3) are generally associated with heterochromatin (condensed DNA, impeding transcription) (*e.g*. (Kimura, 2013);(Cutter DiPiazza et al., 2021)). Limited knowledge is available about acetylated lysine 12 of histone H4 (H4K12ac), mono-methylated lysine 20 of histone H4 (H4K20me1), acetylated lysine 23 of histone H3 (H3K23ac), and tri-methylated lysine 36 of histone H3 (H3K36me3). In humans, H4K12ac is enriched around gene promoters and stretches into gene bodies (Wang et al., 2008); it is also associated with enhancers (Nagarajan et al., 2015). Further studies found that the genes with higher H4K12ac occupancy had a significant bias toward developmental functions during embryogenesis (Paradowska et al., 2012). In cancerous cells, H4K12ac occupancy correlates with gene expression levels (Nagarajan et al., 2015). H4K20me1 is another mark generally associated with transcriptional activation. The most highly transcribed group of human genes tend to have H4K20me1 present (Wang et al., 2008) with strongest enrichment in the first third of the gene body of highly expressed genes, while unexpressed genes are free of H4K20me1 (Segal et al., 2018). However, in combination with H3K9me3, H3K20me1 seems to silence transcription (Sims et al., 2006); this may be because H4K20 mono-methylation is an initial step in the process of generating the highly repressive mark H4K20me3. PR-Set7/Set8/KMT5a is the only enzyme that catalyses mono-methylation of H4K20 and PR-Set7 activity is essential for the earliest stages of mouse embryonic development (Beck et al., 2012). In plants, H3K23ac is localized in gene bodies with an enrichment just downstream of the TSS, but this modification appears to be uncorrelated with gene expression levels (Lu et al., 2015). H3K36 methylation seems associated with actively transcribed chromatin (Georgescu et al., 2020). It has been shown in yeast that H3K36me3 is deposited on histones as they are displaced by RNA polymerase II during transcription. H3K36me3 then serves as a mark for deacetylating histones, preventing probably run-away transcription (Carrozza et al., 2005) (Joshi and Struhl, 2005)). However, H3K36me3 is also used in conjunction with H3K9me3 to repress aberrant transcription (Bartke et al., 2010). H3K36me3 pattern can also influence splicing (Kim et al., 2011), transcriptional repression as well as DNA repair (Sharda and Humphrey, 2022). It was also proposed that H3K36me3 serves to prevent aberrant transcriptional initiation and elongation (Teissandier and Bourc’his, 2017).

In summary, we focused in this study on putatively repressive histone isoforms (H3K9me3, H3K27me3), putatively activating isoforms (H3K4me3, H3K9ac, H4K12ac), and context-depending modifications (H4K20me1, H4K12ac, H3K36me3) that had been linked in the literature to developmental plasticity.

In the present work, we addressed two key questions: (i) what combinations of histone modifications occur around genomic features in *S. mansoni* (it’s “histone code”) and (ii) what changes occur during maturation of the gonads. Answering these questions would contribute to our understanding of the complex male-female interaction, the epigenetic mechanisms involved and the molecular consequences of pairing for female sexual maturation. Prospectively, knowledge in this filed may also provide clues for novel therapeutic options.

We performed ChIP-Seq on isolated ovaries and testes of pairing-inexperienced and pairing-experienced *S. mansoni* worms from single-sex or bi-sex infections using antibodies against the abovementioned histone modifications. We identified those modifications that showed strong changes around genes and repetitive sequences. We subsequently used Hidden-Markov-Models and Random-Forest based techniques to link gene expression to histone modifications. We show here, for the first time in flatworms, (i) combinations of histone modifications (“chromatin colours”) and their putative roles in regulating gene expression, (ii) specific changes in histone modifications in gonads after pairing, and (iii) evidence for epigenetic processes contributing to male-dependent gonad differentiation in the female. Our data additionally supports he concept that histone-modifying enzymes might be potential targets for pharmacological intervention (Pierce et al., 2012; Anderson et al., 2017; Lobo-Silva et al., 2020; Padalino et al., 2023). Our findings also suggest that so far unknown and potentially *Schistosoma*-specific histone modifications exist.

## Results

### Chapter I: The histone code of *S. mansoni* gonads

#### Histone modifications are different around H3K4me3-positive and H3K4me3-negative protein-coding genes

We commenced by analysing the genome-wide coverage of the eight PTMs individually. To this end, we started with chromatin isolated from gonads (testes and ovaries), which had been extracted by an organ isolation approach of pairing-experienced males (bM, bi-sex males), pairing-inexperienced males (sM, single-sex males), pairing-experienced females (bF, bi-sex mature females) and pairing-inexperienced females (sF, single-sex immature females) of *S. mansoni*. In the following we will use the abbreviations: sO (ovaries of sF), bO (ovaries of bF), sT (testes of sM), and bT (testes of bM).

H3K4me3 is an important mark of TSS in many biological models, which motivated us to focus our attention on this mark. Surprisingly, our analyses revealed that there are roughly 40% of all genes that do not possess the H3K4me3 mark while being expressed (Tables 1a and 1b, based on Supplementary file 1).

**Table 1a.** Number of genes with H3K4me3 spanning −2kb to +200 bp of TSS of protein-coding genes (H3K4me3-positives) in the *S. mansoni* gonads. Testes of pairing-experienced males (bT), pairing-inexperienced males (sT), pairing-experienced females (bO) and pairing-inexperienced females (sO) of *S. mansoni*.

|  | sO | bO | sT | bT |
| --- | --- | --- | --- | --- |
| All genes | 5,714<br>(56%) | 5,717<br>(56%) | 5,643<br>(55%) | 5,630<br>(55%) |
| Expressed<br>(>0 rpkm) | 5,695<br>(60%) | 5,697<br>(60%) | 5,624<br>(58%) | 5,609<br>(60%) |

**Table 1b.** Number of genes without H3K4me3 around protein coding genes (H3K4me3-negatives) in the *S. mansoni* gonads.

|  | sO | bO | sT | bT |
| --- | --- | --- | --- | --- |
| All genes | 4,458<br>(44%) | 4,455<br>(44%) | 4,529<br>(45%) | 4,542<br>(45%) |
| Expressed<br>(>0 rpkm) | 3,774<br>(39%) | 3,736<br>(39%) | 4,046<br>(42%) | 3,767<br>(39%) |

Interestingly, most of these genes are common among all gonad stages: (5,599 H3K4me3-positive expressed genes for sT and bT, 5,681 for sO and bO, and 5,472 for all stages; 3,661 H3K4me3-negative genes for sT and bT, 3,525 for sO and bO, and 3,166 for all stages) indicating that almost half of the genes possess no canonical H3K4me3 mark, which has been considered to be typical for active genes in many other organisms. Therefore, we next investigated links between gene function and H3K4me3 enrichment in *cis*.

It had been shown *e.g*. in T-cells that the presence of H3K4me3 is typical for inducible genes (Lim et al., 2009), and it has been proposed that H3K4me3 is a tag constitutively expressed *vs* inducible genes (Murray et al., 2019). However, based on RNA-Seq data (Lu et al., 2016), we observed no bias in the absence of H3K4me3 towards genes that change expression in *S. mansoni* gonad differentiation (Table 2).

**Table 2:** Number and proportion of H3K4me3-negative genes among genes that are up- and downregulated in ovary and testes during transformation from single-sex to sex-experienced gonads. Abbreviations as in table 1.

|  | Down in<br>bO | Up in<br>bO | Down in<br>bT | Up in<br>bT |
| --- | --- | --- | --- | --- |
| Differentially<br>expressed<br>genes | 2,384 | 1,880 | 122 | 97 |
| H3K4me3-negatives in<br>differentially<br>expressed<br>genes | 1,172<br>(49%) | 420<br>(22%) | 79 (65%) | 61<br>(63%) |

The detailed count tables of this are in supplementary file 1.

It could also be that the TSS in H3K4me3-negative genes is erroneously positioned because the mRNA undergoes trans-splicing. Splice leader (SL) trans-splicing is a specialized form of trans-splicing in which a short, capped splice leader exon, derived from a distinct small splice leader RNA, is transferred to the 5’end of a separate pre-mRNA. This reaction replaces the original 5’end of the pre-mRNA and produces a mature mRNA with a uniform leader sequence. For that, we analysed 4,754 trans-spliced genes (Boroni et al., 2018) and compared it to the H3K4me3-neg genes. In sO 796 matched, in bO 795, in sT 805 and 812 in bT, *i.e*. less than 17% of all trans-spliced genes are free of H3K4me3 meaning that SL trans-splicing is unlikely to be the main origin of this observation.

We then investigated the difference in abundance of mRNA between H3K4me3-positive and H3K4me3-negative genes. We found that H3K4me3-positive genes have, in average, a 5.8-fold increase in log(RPKM) (Reads Per Kilobase of transcript per Million mapped reads) compared to negative genes, regardless of whether summarized over all four sample types or analysed by sex or maturation/differentiation state. The differences are highly significant (Figure 1).

**Figure 1.**
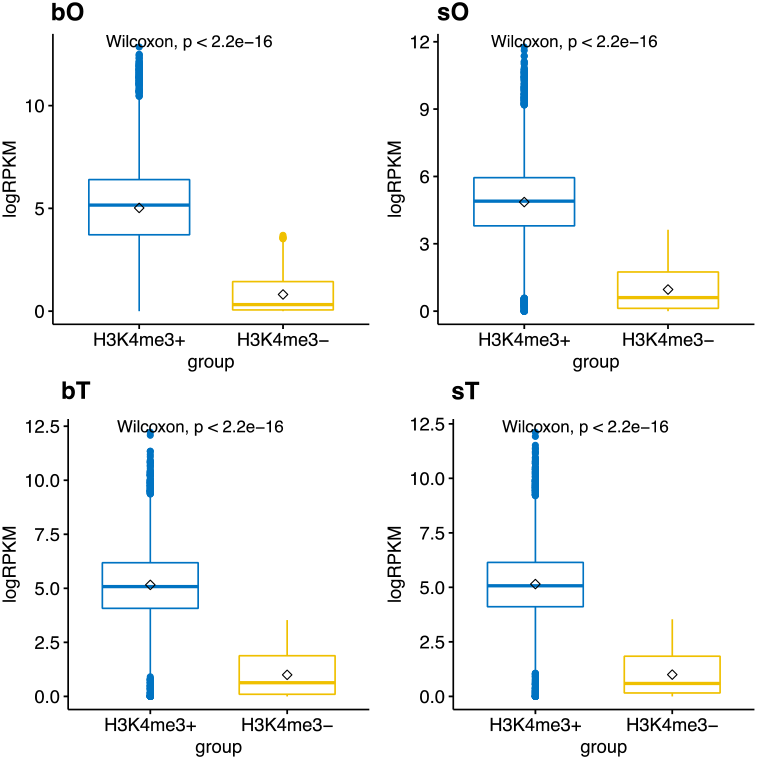
Boxplots and results of Wilcoxon test of RNA-Seq data and H3K4me3 presence in *cis*. Left/Blue: H3K4me3-positive genes; right/yellow: H3K4me3-negative genes; y-axis log(RPKM). Number of genes as in tables 1a and 1b.

To further investigate if there was any functional significance in the absence of H3K4me3, we employed Rank Based Gene Ontology Analysis using H3K4me3 count values per gene in supplementary file 1 as the input and gene GO association from WormBaseParasite (supplementary file 2). Results are summarized in supplementary file 3. In brief, H3K4me3-negative genes were associated with signal transduction, cell membrane, and extracellular matrix as well as RNA production and protein secretion *i.e*. interaction with the extracellular space. In contrast, H3K4me-positive genes were associated with catabolic and metabolic activities and the inner membrane systems including nuclear envelope and the nucleus *i.e*. the “inner” world of the cell. The presence of H3K4me3-expressed genes is indicative for non-canonical chromatin structure in schistosome gonads *i.e*. not shared between mammalian host and parasite, and that chromatin structure is used by the schistosome to control the gene functions in the gonads, probably in a reciprocal, pairing-dependent manner.

In summary, H3K4me3 presence around the TSS of a gene is indicative for higher transcription and is associated with GO terms that do not relate to the interaction with the environment, but it is not suggestive for inducible or uninducible genes.

#### Individual histone modifications show specific enrichment around H3K4me3-positive and -negative protein-coding genes, and genes encoding lncRNA

Hierarchical clustering by metagene profiles of all histone modifications, except H3K4me3, confirmed 3 types of genes (H3K4me3-positive genes, H3K4me3-negative genes, and lncRNA genes), and each group was investigated (Figure 2) separately. Of note, H3K4me3-negative genes in ovaries of sF were identified as a clear outlier.

**Figure 2.**
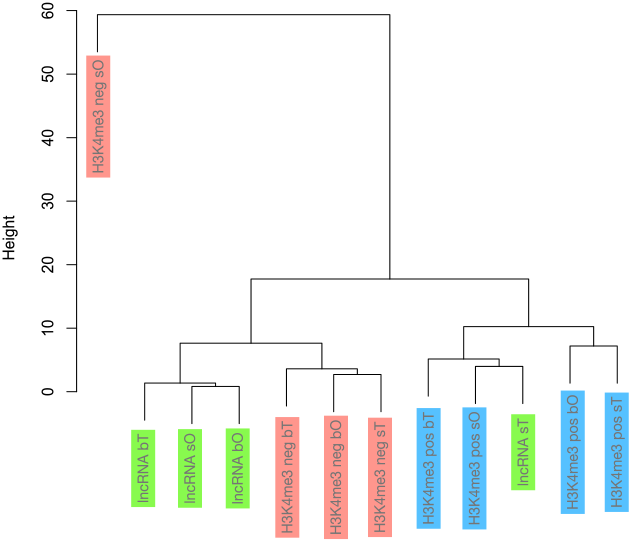
Clustering into 3 types of genes by combining seven ChIP metagene profiles (excluding H3K4me3 profiles that were already used to differentiate H3K4me3 positive and negative genes).

**Figure 3a.**
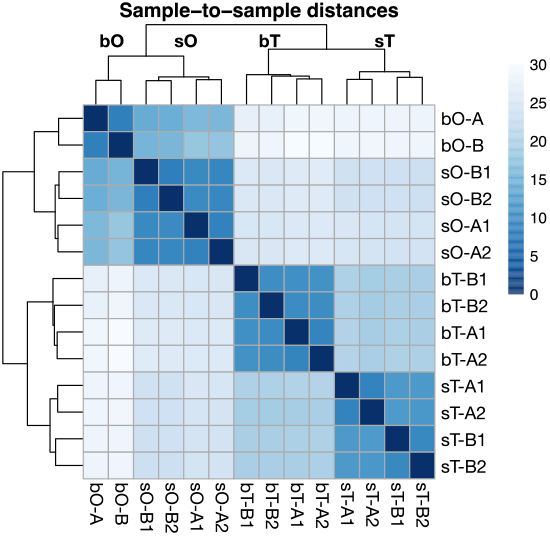
Heatmap of H3K23ac changes in repetitive sequences. Shades of blue represent Euclidian distances. A and B are biological replicates, 1 and 2 technical replicates. Clustering clearly distinguishes between the four samples types (bO, sO, bT, sT).

**Figure 3b.**
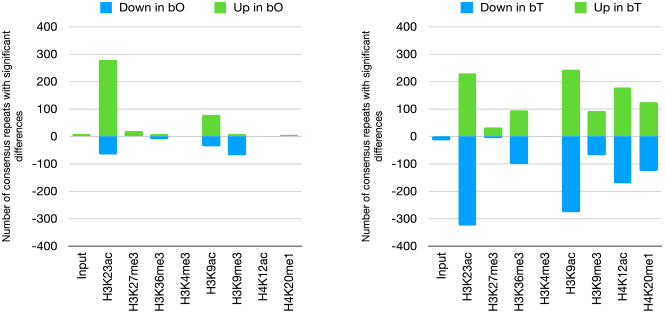
Changes in histone modifications in repetitive sequences.

#### Individual histone modifications in repetitive sequences

Repetitive sequences are now recognised as an important component of the genome. In *S. mansoni*, repeats are transcribed in stage specific manner (Lepesant et al., 2012). We, therefore, wondered if repeats would show differential histone modifications in gonads. Alignment of short-read sequences as in ChIP-seq cannot distinguish between different copies of repeats along the genome. Consequently, we resorted to a type of analysis in which we used a set of 3,145 repeat consensus sequences that had been previously produced (Lepesant et al., 2012). ChIP-Seq reads were aligned to these consensus sequences and hits were counted. This averaging approach provides a global view of histone modifications in repetitive sequences and indicates the complete absence of H3K4me3 in these sequences in both sexes. Interestingly, H3K23ac is highly enriched in repeats in both ovaries and testes, and shows strong differential enrichment between the sexes, but also between mating inexperienced and experienced gonads. Other important differences between sT and bT occur in H3K9ac, H4K12ac, and H4K20me1 (Figure 3 and supplementary file 4).

At this stage of the analysis, where histone modifications are examined individually, and all analyses are performed in parallel, we define four categories of features, each “painted” by a specific histone modification: protein-coding genes with or without H3K4me3 enrichment at the TSS, lncRNA genes without clear enrichment (for details see (Augusto et al., 2025), and repetitive sequences strongly marked by and divergent in H3K23ac. The strong sex- and developmental specific plasticity of chromatin around repeats is particularly interesting and warrants further investigation that is beyond the scope of the current work, which focussed on protein coding genes.

#### Chromatin of S. mansoni gonads has 54 predominant colours

Genome-wide analyses of testes using ChromstaR uncovered 248 of 256 possible chromatin colours, and in ovaries we found all 256 possible histone combinations. However, we also realised that some of the colours occurred very rarely, while others were abundant. To our knowledge, there is no consensus on how to evaluate the biological importance of a chromatin colour. We reasoned that a colour would likely be biologically important if it covers a reasonable amount of the genome, and if it is present in the regulatory regions of genes. Therefore, we arbitrarily established the following criteria for biologically significant chromatin colours: in any of the four states of gonads (i), it covers at least 1% of the genome, (ii) it covers at least 0.5% of the total number of TSSs, and (iii) it is among the top 10 colours that can be associated with gene expression.

Based on criteria (i) and (ii), we identified 34 colours (supplementary file 5). The chromatin colours seen in sT are mostly enriched in marks such as H4K12ac, H3K23ac, and H3K27me3. In bT, the most enriched chromatin colours included H3K9ac and H3K9me3. In sO, around 10% of the genome is decorated with the chromatin colour [H3K23ac+H3K27me3+H3K9ac+H4K12ac+H4K20me1]. In bO, nine colours were selected based on our criteria, but the most abundant chromatin colour was [H3K23ac+H3K27me3+H3K36me3+H3K9ac+H4K12ac+ H4K20me1], covering more than 20% of the entire genome. Knowing that the levels of histone modifications are well correlated to gene expression in other species, and assuming that this relationship can be generalized across different *S. mansoni* gonad conditions, we then performed a random forest analysis to correlate RNA-Seq and ChIP-Seq data (criterion iii). We selected the top 10 most important colours for gene expression giving a total of 28 colours from all 4 conditions (supplementary file 5). Unsurprisingly, most of theoretically possible colours had not enough reads counts to affect the correlation to gene expression, which reinforces the idea that not all colours are equally important for gene expression (supplementary file 6).

Combining the list of chromatin colours fulfilling criteria (i-ii) with the list corresponding to criteria (iii) we established a list of 54 non-redundant chromatin colours that we call here “biologically significant” colours (supplementary file 5).

#### Chromatin colours are associated with genome features

In previous studies on other biological models, it was evident that specific chromatin colours occur in “active” or “repressive” or “Polycomb” regions of the genomes (Filion et al., 2010). To investigate the possible association of predominant chromatin colours with annotation features of the *S. mansoni* genome we applied the clustering algorithm of ChromstaR. Unsupervised clustering of genomic features with the *plotFoldEnrichHeatmap* function of ChromstaR clearly clustered features by colour similarity into 3 groups in each of the 4 samples (sO, bO, sT, and bT): (i) 2 kb upstream (promoter), 5’UTR, exon, ATAC-Seq-positives in males and females, (ii) gene, mRNA, lncRNA, 3’UTR and ATAC-seq negatives in male and female, and (iii) pseudogenes (Figure 4).

**Figure 4.**
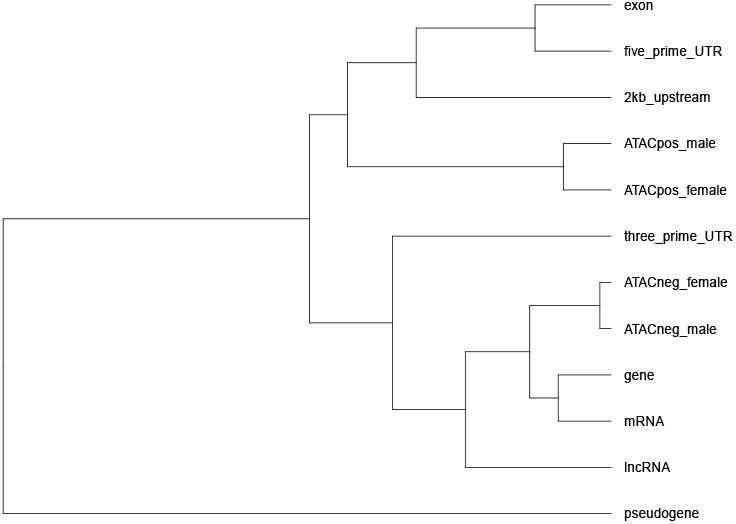
Clustering of features by significant colour similarities (example for mature testis). Exons and putative promotor regions show similar histone modifications and are associated with Tn5 accessible chromatin (ATACpos_male and ATACpos_female). Gene bodies, regardless of if they code for lncRNA or mRNA, have slightly different histone modification combinations from the former. Pseudogenes are clearly painted with very different histone modifications. Clustering results for sO, bO, and sT in supplementary files 7.

However, there was not a single specific colour that painted these features. In other words, *e.g*. the feature “exon” is characterised by the enrichment of 10 colours, the composition of which was similar to the composition spanning the feature “2 kb upstream” but definitely different from those in the feature “pseudogene”. This indicates a complex and ‘multi-colour’ chromatin, but nevertheless regular painting of genome features with histone modifications. Details are given in supplementary file 7.

#### Chromatin colours occur collinearly with transcription steps along protein coding genes

To gain insights into the link between proximal histone modifications and transcription, we aggregated the colour profiles by unsupervised hierarchical clustering into groups that showed similar metagene profiles based on log(observed/expected) values within and ±2kb around genes. We allowed up to 6 clusters, and we identified finally 2-4 types per sample. Initial clustering was done for all 265 chromatin colours; a more detailed analysis of the metagene profiles was done only for the 54 predominant colours.

In H3K4me3-positive genes of ovaries, there are 3 modification types: (i) upstream of the TSS, and presumably pre-initiation complex recruitment-associated patterns are characterized by H3K27me3 and H3K9me3 in combination with other marks. Pattern (ii) peaks around the TSS and consistently featuring H3K4me3 alone, or in combinations, suggesting an association with promoter relaxation. Pattern (iii) is complex but lacks H3K9me3 and H3K4me3, and it is possibly associated with transcription elongation and termination (Figure 5a).

**Figure 5a.**
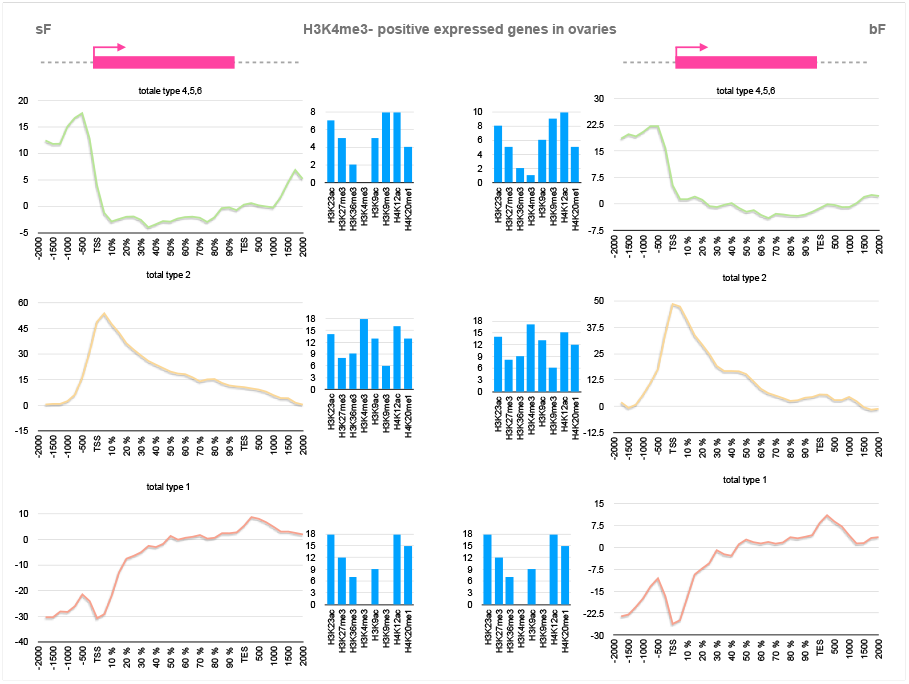
Metagene profiles of cumulated log(obs/exp) values for profiles that were clustered by similar shapes in the group of H3K4me3-positive expressed protein-coding genes in ovaries. sO (left) and bO (right). Y-axes, sums of log(obs/exp). The composition of the underlying colours is included in the supplementary data. Histograms in the middle show occurrence of each histone modification in the colours.

**Figure 5b.**
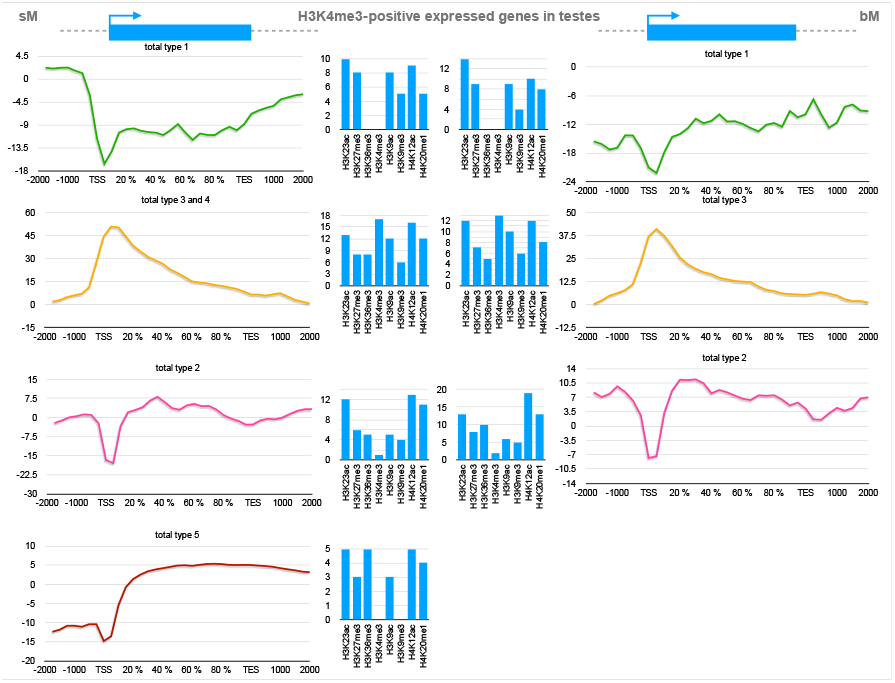
Metagene profiles of cumulated log(obs/exp) values for profiles that were clustered by similar shapes in the group of H3K4me3-positive expressed protein coding genes in testes. sT (left) and bT (right).

**Figure 5c-d.**
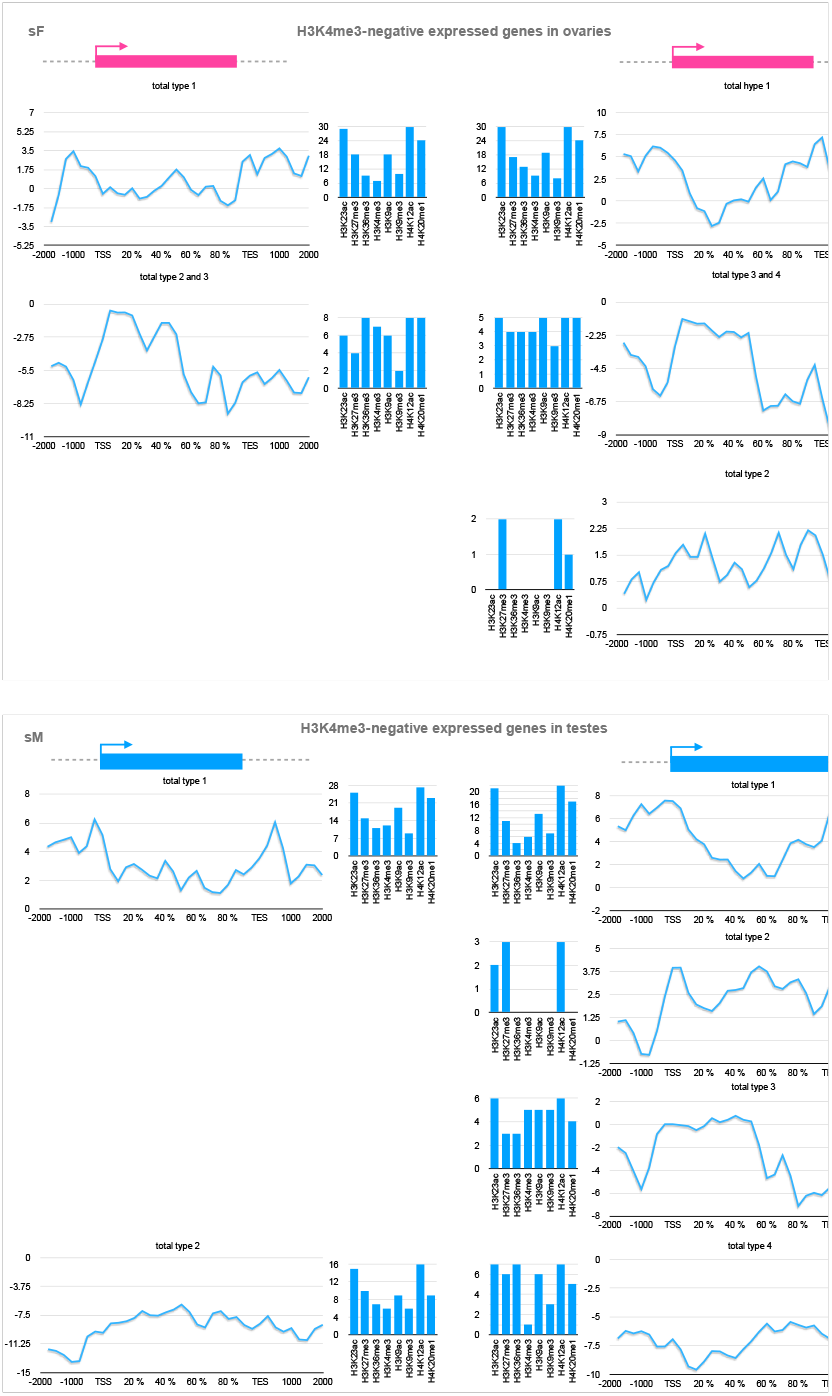
Metagene profiles of cumulated log(obs/exp) values for profiles that were clustered by similar shapes in the group of H3K4me3-negative expressed protein coding genes in ovaries (upper panel) and testes (lower panel). sO (top left), bO (top right) and sT (bottom left), bT (bottom right).

In H3K4me3-positive genes of testes, there are up to 4 modification types: (i) a sT pattern that is enriched in H3K27me3 and H3K9ac upstream of the TSS and. This enrichment was absent from bT. (ii) Similar to the sO/bO, sT/bT show a pattern with a strong enrichment of H3K4me3 around the TSS. (iii) The third pattern spans the gene bodies homogeneously with strong H4K12ac marks. Finally (iv), a pattern that is only present in sT shows strong enrichment of H3K36me3 downstream of the TSS and might be associated with transcription elongation (Figure 5b).

Conversely, H3K4me3-negative genes exhibit patterns that cannot easily be associated with the different stages of transcription (Figure 5c-d). Details are in supplementary file 7.

#### H3/H4 acetylation is best predictor for gene expression in H3K4me3-positive protein coding genes

Collinearity of histone modifications and transcription stages are strongly suggestive for a role in control of transcription. We, therefore, hypothesized that individual histone modifications or chromatin colours were associated with mRNA presence of protein coding genes *i.e*. that some colours are predictive for gene expression. Unsurprisingly, when all genes were considered, H3K4me3 presence around the TSS was the best predictor for gene expression. But this is due, as shown above, to the low expression in the 40% of the expressed genes that do not possess H3K4me3. If studied apart, random forest performs best with a training set of bF, but the correlation between prediction and measured RPKM is only around 0.5 for all samples. H4K12ac, H3K23ac and H3K9ac are the most important predictive modifications for mRNA abundance. This means if genes carry a H3K4me3 mark around their TSS, their transcription status is positively associated with histone acetylation.

#### H3/H4 acetylation and H3K36me3 are best predictors for gene expression in H3K4me3-negative protein coding genes in immature gonads

We used the same approach as above for genes that did not contain H3K4me3 counts around the TSS (Table 1b). We found much weaker correlation between histone modifications and RNA-Seq data than for H3K4me3-positive genes (0.34 for bOF, 0.47 for sO, 0.55 for bT, and 0.61 for sT). The main predictors of gene expression are H3K9ac and H3K36me3 in sO, H3K9ac, and H4K12ac in bO, H4K12ac and H3K36me3 in sT, and H4K12ac and H3K9ac in bT. Noteworthy is the loss of importance of H3K36me3 during gonad maturation of ovaries, which also occurs, albeit much less degree, in transition from sT to bT. Details are in supplementary file 7.

### Chapter II: Histone modifications in the gonads after pairing

#### Chromatin of S. mansoni gonads increases in complexity after paring

To describe the genome-wide coverage of the eight studied PTMs in bM, sM, bF, and sF gonads, we analysed the frequency of each histone modification at given genomic position.

In general, in gonads of paring-experienced worms, the occurrence frequency of the most studied histone modifications increases compared to pairing-inexperienced worms (bT vs sT; bO vs sO). In testes of bM, the coverage of H4K12ac, H3K9ac and H3K23ac increased by 286, 5,695, and 862-fold in comparison to sM (Table 3a). In the bF ovaries, the coverage of H3K36me3, H3K9ac, and H3K23ac increased by 217, 261 and 66-fold in comparison to the ovaries of sF (Table 3b).

**Table 3a.** Proportion of the genome in the testis of adult *S. mansoni* pairing-experienced males (bT) and pairing-inexperienced males (sT) that is painted with H3K4me3, H3K27me3, H3K9me3, H3K9ac, H3K23ac, H4K12ac, H4K20me1 and H3K36me3. Column sM+bM indicates proportion in which the PTM occurs in both samples, column “Absence” the proportion of the genome that is free from the PTM.

|  | sT-specific | bT-specific | sT+bT | Fold change |
| --- | --- | --- | --- | --- |
| H3K4me3 | 0.04% | 0.01% | 3.48% | 0.32 |
| H3K27me3 | 0.64% | 5.19% | 58.53% | 8.07 |
| H3K9me3 | 0.57% | 50.52% | 11.87% | 89.19 |
| H3K9ac | 0.00% | 19.82% | 50.29% | 5695.19 |
| H3K23ac | 0.01% | 7.16% | 69.40% | 862.04 |
| H4K12ac | 0.01% | 3.17% | 71.67% | 286.62 |
| H4K20me1 | 0.14% | 16.05% | 38.14% | 116.29 |
| H3K36me3 | 1.72% | 10.34% | 19.44% | 6.00 |
| Average | 0.39% | 14.03% | 40.35% |  |

**Table 3b.** As for 1a but here ovaries of pairing-experienced females (bO), and pairing-inexperienced females (sO).

|  | sO-specific | bO-specific | sO+bO | Fold change |
| --- | --- | --- | --- | --- |
| H3K4me3 | 0.02% | 0.04% | 4.03% | 1.90 |
| H3K27me3 | 0.54% | 3.01% | 50.94% | 5.56 |
| H3K9me3 | 0.13% | 0.13% | 5.77% | 0.95 |
| H3K9ac | 0.03% | 8.46% | 49.64% | 261.02 |
| H3K23ac | 0.08% | 5.47% | 56.11% | 66.07 |
| H4K12ac | 0.37% | 0.22% | 58.72% | 0.61 |
| H4K20me1 | 1.08% | 0.11% | 61.68% | 0.10 |
| H3K36me3 | 0.07% | 15.10% | 46.71% | 217.78 |
| Average | 0.29% | 4.07% | 41.70% |  |

With respect to the number of peaks of each histone modification, genome-wide analyses also revealed an increase in the gonads from paring-inexperienced (sT and sO) to paring-experienced (bT and bO) worms. In male gonads, H4K20me1 was the most abundant PTM with 1,246 peaks exclusively in sT, 72,228 peaks exclusively in bT, and 12,1621 in both conditions. Regarding the sT vs bT samples, acetylated histones H3K9ac, H3K23ac, H4K12ac were highly variable in male gonads (Table 4a). Paring also promoted chromatin modifications in the ovary. H3K27me3 was the most abundant PTM with 43,184 peaks in total (5,340 peaks in sO, 16,527 peaks in bO, and 48,795 in both conditions). H3K36me3 and acetylation of H3K9 and H3K23 were the most enriched histone modification from sO to bO transition (Table 4b).

**Table 4a.** Number of peaks along the genome in the testis of adult *S. mansoni* pairing-experienced males (bT) and pairing-inexperienced males (sT) for H3K4me3, H3K27me3, H3K9me3, H3K9ac, H3K23ac, H4K12ac, H4K20me1 and H3K36me3.

|  | sT-specific | bT-specific | sT+bT | Fold change |
| --- | --- | --- | --- | --- |
| H3K4me3 | 370 | 47 | 6 384 | 0.13 |
| H3K27me3 | 5 271 | 27 093 | 60 931 | 5.14 |
| H3K9me3 | 5 834 | 71 841 | 31 563 | 12.31 |
| H3K9ac | 23 | 82 558 | 82 160 | 3589.48 |
| H3K23ac | 59 | 28 307 | 29 798 | 479.78 |
| H4K12ac | 70 | 19 783 | 28 790 | 282.61 |
| H4K20me1 | 1 246 | 72 228 | 12 1621 | 57.97 |
| H3K36me3 | 4 872 | 9 198 | 10 620 | 1.91 |

**Table 4b.** As for 2a but here ovaries of pairing-experienced females (bO) and pairing-inexperienced females (sO).

|  | sO-specific | bO-specific | sO+bO | Fold change |
| --- | --- | --- | --- | --- |
| H3K4me3 | 165 | 167 | 6 780 | 1.01 |
| H3K27me3 | 5 340 | 16 527 | 48 795 | 3.10 |
| H3K9me3 | 380 | 1 177 | 5 558 | 3.10 |
| H3K9ac | 363 | 39 761 | 40 286 | 109.53 |
| H3K23ac | 891 | 23 885 | 41 199 | 26.81 |
| H4K12ac | 4 143 | 2 557 | 42 820 | 0.62 |
| H4K20me1 | 10 853 | 1 346 | 42 980 | 0.12 |
| H3K36me3 | 744 | 87 647 | 88 817 | 117.80 |

Highlighting the importance of histone acetylation activity in the transition of single-sex to bi-sex testes and ovaries.

#### Specific chromatin colours change in ovaries upon pairing

Previously, we had used RNAseq to identify 3,600 genes that change in expression during ovary maturation, of which 1,752 genes were found to be upregulated and 1,848 downregulated in bO compared to sO (Lu et al., 2016). For convenience, we will refer to the top 100 most upregulated genes and top 100 most downregulated genes during ovary maturation as the *ovary-upregulated* and *ovary-downregulated* sets, respectively. We compared the overlaps between *ovary-upregulated* genes (including their UTRs) with chromatin colour blocks in sO, as well as their overlaps with colour blocks in bO. We found overlaps with colour blocks in bO being enriched for colours involving H3K36me3, compared to the overlaps with colour blocks in sO (supplementary file 8.1). The same trend was also observed for downregulated genes: their overlaps with chromatin colour blocks in bO were enriched for colours involving H3K36me3, compared to overlaps with colour blocks in sO (supplementary file 8.2). This suggested that genes upregulated or downregulated during ovary maturation seem to be more decorated with colours involving H3K36me3 in bO compared to sO. However, for both the *ovary-upregulated* and *ovary-downregulated* genes, the chromatin colours that changed most in these genes during ovary maturation did not involve H3K36me3 but [H3K23ac+H3K27me3+H3K9ac+H4K12ac+H4K20me1]. That is, 19.8% (20.1%) of the overlaps between *ovary-upregulated* (*ovary-downregulated*) genes with colour blocks in sO showed this colour, but this reduces to 4.8% (4.2%) of overlaps with colour blocks in bO (supplementary files 8.1 and 8.2)).

Next, we compared a set of 905 genes identified by (Lu et al., 2016) as up-regulated in all gonads (testis and ovary of paired and unpaired males and females, respectively) to those of whole worms with ≥1.5-fold difference (1,012 genes in the original paper; mapped to 905 current identifiers). We will refer to these as *gonad-enriched* genes. The *gonad-enriched* genes are expressed in ovaries/testes but their expression does not change as much during ovary maturation as the *ovary-upregulated*/*ovary-downregulated* genes. Overlaps between the gonad-enriched genes and the chromatin colour blocks in bO appeared enriched for colours containing H3K9ac, compared to overlaps with chromatin colour blocks in sO (supplementary file 8.3), although the trend was not as striking as that seen for H3K36me3 and the *ovary-upregulated*/*ovary-downregulated* genes. As for the *ovary-upregulated*/*ovary-downregulated* genes, the colour that changed most in *gonad-enriched* genes during ovary maturation was [H3K23ac+H3K27me3+H3K9ac+H4K12ac+H4K20me1], although it did not reduce as much during ovary maturation as for the *ovary-upregulated*/*ovary-downregulated* genes. We found this colour in 7.5% of the overlaps between *gonad-enriched* genes and colour blocks in sO, compared to 2.8% of overlaps in bO (supplementary file 8.3).

The trends described above were found by taking a particular set of genes (e.g. *ovary-upregulated*) and looking at all their overlaps with chromatin colour blocks in sO compared to their overlaps to colour blocks in bO. However, we were not sure whether these trends resulted from an addition of particular marks (*e.g*. H3K36me3) to existing blocks or from the creation of totally new colour blocks during ovary maturation. To investigate this, we identified colour blocks in sO that had the exact same coordinates (start and end) as a colour block in bO, or nearly the same coordinates (covering ≥90% of the block in bO, and ≥90% of the overlapping block in sO). We call these *position-stable colour blocks*. Taking the *position-stable colour blocks* within *ovary-upregulated genes* (and their UTRs), 56.6% (146/258) of those that differed in colour, differed by addition of H3K36me3 in the mature ovaries (supplementary file 8.4). For example, such a change could be a [H3K9ac+H3K23ac] colour block in sO being replaced by an exactly overlapping [H3K9ac+H3K23ac+H3K36me3] colour block in bO. The second most common change was +H3K9ac. Likewise, for *position-stable colour blocks* within *ovary-downregulated* genes, we found 58.8% (133/226) of the blocks that differed in colour, also differed by addition of H3K36me3 in bO (supplementary file 8.4). The second most common difference was +H3K9ac. In both sets of genes, the addition of H3K36me3 was the most common difference in chromatin colour in *position-stable colour blocks*.

In gonad-enriched genes, the most frequent chromatin change among *position-stable colour blocks* was the acquisition of H3K9ac, representing 27.3% (942/3,446) of all blocks showing a colour change (Supplementary File 8.4). The second most common change was the acquisition of H3K36me3, accounting for 24.7% (851/3,446) of the differing blocks. This proportion was significantly lower than that observed for H3K36me3 gain in *position-stable colour blocks* of *ovary-upregulated genes* and *ovary-downregulated genes* (Fisher’s exact test, P = 10^-15^ for both comparisons).

The findings above suggested that the chromatin in *ovary-upregulated*/*ovary-downregulated* genes tend to be more highly decorated with H3K36me3 in bO than in sO, but that *gonad-enriched* genes showed far less difference in H3K36me3 marks between immature and mature ovaries. It is clear that more H3K36me3 is added to *ovary-upregulated*/*ovary-downregulated* genes than to *gonad-enriched* genes during ovary maturation. We next investigated whether, before ovary maturation, *ovary-upregulated*/*ovary-downregulated* genes start off in sO with a lower level of H3K36me3 compared to *gonad-enriched* genes. Taking chromatin colour blocks in sO, overlaps between these blocks and *ovary-upregulated* genes were enriched for colours lacking H3K36me3 but having H3K9ac marks, compared to overlaps between these blocks and *gonad-enriched* genes (supplementary file 8.5). As mentioned above, 19.8% of overlaps between *ovary-upregulated* genes and colour blocks in sO were due to [H3K23ac+H3K27me3+H3K9ac+H4K12ac+H4K20me1], but this colour was only responsible for 7.5% of overlaps between *gonad-enriched* genes and colour blocks in sO (supplementary file 8.5).

In repetitive sequences, we observed little changes in histone modifications except for acetylation in H3K23 that increased in almost 300 consensus repeats and decreased in roughly 50, from sO to bO (Figure 5 and supplementary data 4).

#### Specific chromatin colours change in testes upon paring

In our previous RNAseq study, we identified 243 genes that changed in expression between sM and bM, 96 being upregulated and 147 downregulated [Lu et al 2016 PMID:27499125]. We will refer to the 96 upregulated genes as *testis-upregulated* and the top 100 most downregulated genes as *testis-downregulated*. We found overlaps of *testis-upregulated* genes with colour blocks in bT, which were enriched for colours involving H3K9me3, compared to overlaps with colour blocks in sT (supplementary file 8.6); the same trend was also seen for *testis-downregulated* genes (supplementary file 8.7). Thus, genes upregulated/downregulated in testes tend to have less H3K9me3 in sT compared to bT. Overlaps between the *gonad-enriched* genes and the chromatin colour blocks in bT also appeared enriched for colours containing H3K9me3 (supplementary file 8.8), although the differences between sT and bT were not as dramatic as for *testis-upregulated*/*testis-downregulated* genes.

In both the *testis-upregulated* and *testis-downregulate*d genes, the chromatin colour that changed most during testis maturation was [H3K23ac+H3K27me3+H3K9ac+H3K9me3+H4K12ac+H 4K20me1]; this was present in 2.6% (1.8%) of overlaps between *testis-upregulated* (*testis-downregulated*) genes with colour blocks in sT, but increased to 21.8% (20.4%) of overlaps with colour blocks in bT (supplementary files 8.6 and 8.7). In contrast, for *gonad-enriched* genes, this colour changed less, from 0.8% of overlaps with colour blocks in sT to 7.8% of overlaps in bT (supplementary file 8.8). In fact, the colour that changed most in *gonad-enriched* genes was [H3K23ac+H3K27me3+H3K36me3+H3K9ac+H3K9me3+ H4K12ac+H4K20me1], which increased from 2.9% of overlaps in sT to 17.1% of overlaps in bT (supplementary file 8.8).

When we looked at *position-stable colour blocks*, this time considering those that have stable positions in sT and bT, the same patterns emerged, with H3K9me3 being added to both *testis-upregulated*/*testis-downregulated* and *gonad-enriched* genes, but with a greater increase of H3K9me3 in *testis-upregulated*/*testis-downregulated* than *gonad-enriched* genes. That is, for *position-stable colour blocks* that lie within *testis-upregulated* genes (and their UTRs), 59.8% (502/840) of those that differed in colour between sT and bT, differed by addition of H3K9me3 in the bT (supplementary file 8.9). The second most common change was +H3K9ac (supplementary file 8.9). Similarly, for *testis-downregulated* genes, 64.4% (601/933) of the *position-stable colour blocks* that were variable in colour differed by addition of H3K9me3 in bT (supplementary file 8.10). The second most common change was +H3K9ac,+H3K9me3 and the third most common +H3K9ac (supplementary file 8.10).

In contrast, for the *gonad-enriched* genes, even though addition of H3K9me3 was still the most common change in *position-stable colour blocks*, only 43.0% (2335/5428) of the *position-stable colour blocks* that changed in colour differed by addition of H3K9me3 (supplementary file 8.11). The second most common change was +H3K9ac. That is, for *gonad-enriched* genes and for *position-stable colour blocks* that changed in colour between sT and bT, significantly fewer genes changed by addition of H3K9me3 (43.0%) compared to in *testis-upregulated* genes (59.8%; Fisher’s test: *P*=10^-15^) or *testis-downregulated* genes (64.4%; *P*=10^-15^

We next investigated whether *testis-upregulated*/*testis-downregulated* in sT have a lower level of H3K9me3 compared to *gonad-enriched* genes. Considering colour blocks in sT, overlaps between these blocks and *testis-upregulated* genes seem to have similar frequencies of chromatin colours involving H3K9me3, compared to overlaps between these blocks and *gonad-enriched* genes (supplementary file 8.12). Thus, it appears that in single-sex testes, the *testis-upregulated* genes do not differ much in their level of H3K9me3 compared to the *gonad-enriched* genes; therefore, both sets of genes seem to start off with a similar level of H3K9me3 before pairing with a female, but *testis-upregulated* genes seem to gain more H3K9me3 marks than *gonad-enriched* genes following pairing.

Of note, overlaps between the colour blocks in sT and *gonad-upregulated* genes seem to be enriched for colours involving H3K36me3, compared to the overlaps between these blocks and *testis-upregulated* genes (supplementary file 8.12). In other words, *testis-upregulated* genes seem to have lower levels of H3K36me3 than *gonad-enriched* genes in sT. In particular, two chromatin colours lacking H3K36me3 are particularly common in *testis-upregulated* genes, compared to *gonad-enriched* genes, in sT: [H3K23ac+H3K27me3+H3K9ac+H4K12ac] and [H3K23ac+H3K27me3+H3K9ac+H4K12ac+H4K20me1] (supplementary file 8.12). Therefore, as for the *ovary-upregulated* genes, a low level of H3K36me3 in *testis-upregulated* genes in sT may allow changes in expression levels on the way to bT (although, unlike *ovary-upregulated* genes, they do not tend to gain H3K36me3 marks).

A detailed metagene analysis around lncRNA coding genes (Augusto et al., 2025) also revealed that chromatin colours exhibit a tendency to aggregate based on the similarities in their metagene shapes, leading to the formation of less than six distinct clusters. Similarly to what we find in protein coding genes, these clusters can be further grouped according to their resemblances by shape, which are co-linear with specific regions of the genes and potentially associated with transcriptional stages. We also showed that there are strong changes in the composition of the clusters depending on the gonad type.

Concerning repetitive DNA, there were more complex changes than in the ovaries, and this for all investigated histone modifications (except for H3K4me3 that is absent from repeats, see above).

Given the massive histone modification changes, it was unsurprising that a differential histone mark enrichment analysis with ChromstaR identified thousands of differential chromatin colour peaks between gonads of pairing-experienced versus inexperienced worms (94,148 regions in sO vs bO; 249,867 regions in sT vs bT, table 3). This analysis confirmed the predominant role of H3K9me3 in testes, H3K36me3 in ovaries, and H3K9 and H3K23 acetylation in the gonads of both sexes.

**Table 3.** number of differential peaks for individual histone modifications.

|  | H3K23ac | H3K27me3 | H3K36me3 | H3K4me3 | H3K9ac | H3K9me3 | H4K12ac | H4K20me1 |
| --- | --- | --- | --- | --- | --- | --- | --- | --- |
| sT-not-bT | 11 | 659 | 693 | 87 | 2 | 11 | 9 | 163 |
| bT-not-sT | 13,118 | 475 | 930 | 12 | 17,790 | 33,407 | 4,427 | 297 |
| sO-not-bO | 8 | 184 | 10 | 19 | 3 | 2 | 787 | 2,851 |
| bO-not-sO | 11,011 | 2,441 | 32,821 | 88 | 11,165 | 102 | 354 | 139 |

We, therefore, performed combinational metagene ChromstaR analyses with H3K23ac, H3K9ac, and H3K36me3, in which major global changes occurred in ovaries, and H3K23ac, H3K9ac, H4K12ac, and H3K9me3 in testes. Results for the chromatin colours that changed around genes during maturation are summarized in figures 6a and 6b.

**Figure 6a.**
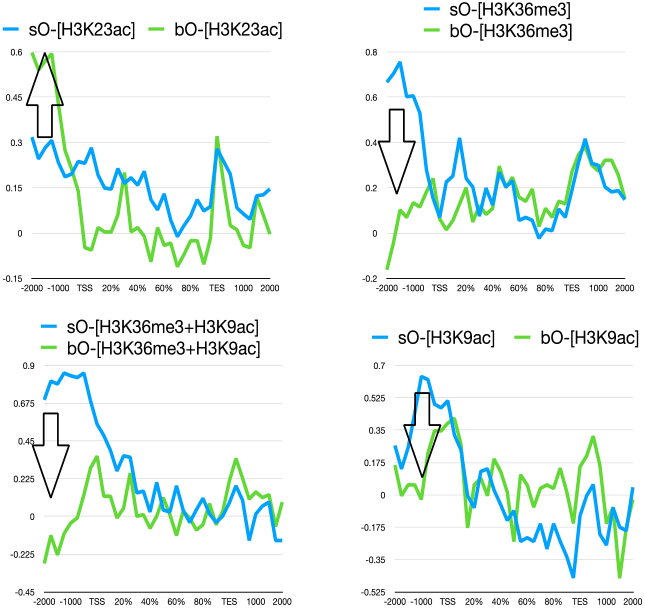
Metagene profiles of ovaries in a subset of chromatin colours, for which changes occurred around genes during the maturation of the ovaries. Blue: sO, green: bO. Arrow indicates direction and location of change from sO to bO.

**Figure 6b.**
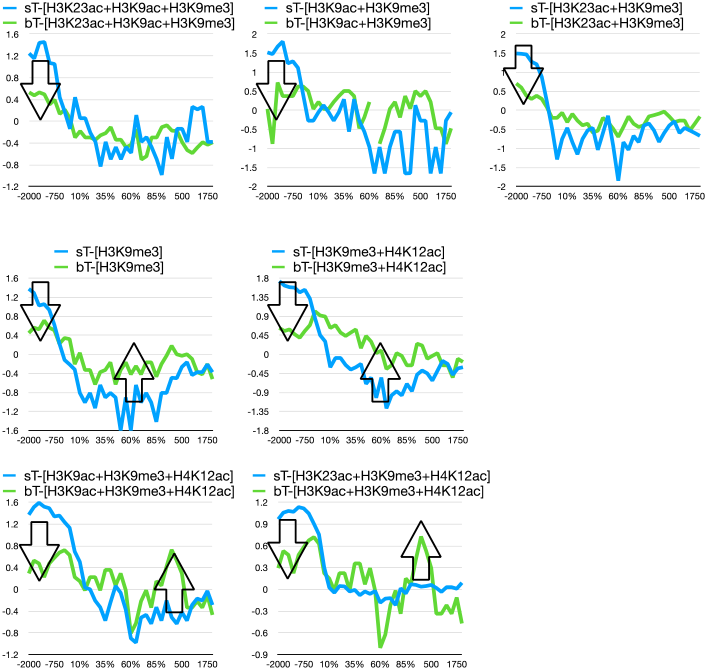
Metagene profiles of testes in a subset of chromatin colours for which changes occurred around genes in the maturation of the testes. Blue: sT, green: bT. Arrows indicate the direction and location of changes from sF to bF.

In conclusion, all our analyses converged towards a particular importance of histone acetylation and H3K9/H3K36 trimethylation in gonads from single-sex to paired worms. Histone acetylases and deacetylases are known not to be very specific for their (de)acetylation sites. Therefore, we focused our attention on enzymes involved in the methylation and de-methylation pathways of these histone PTMs.

### Homology searches identify genes for H3K9/H3K36-specific methyltransferases and demethylating enzymes in S. mansoni

H3K36me3 is an evolutionary conserved modification, and enzymes responsible for H3K36 trimethylation have been identified in several model organisms. Homology searches against *Caenorhabditis elegans, Drosophila melanogaster*, and Human enzymes identified Smp_210650 as the most likely *S. mansoni* SETD2 H3K9/H3K36-specific methyltransferase ortholog. Based on the gene expression atlas (https://v7test.schisto.xyz), Smp_210650 shows highest expression in ovaries and a log2fold-overexpression of 2.7 in bO (http://schisto.xyz). In contrast, adult worm single cell data in https://www.collinslab.org/schistocyte/clusters.html indicate higher expression of Smp_210650 in male neoblasts. No phenotype was observed by knock-down (Wang et al., 2020). Smp_133910, which had earlier been identified as SETD2 candidate (Cabezas-Cruz et al., 2014), is now merged with Smp_210650 in v7 of the genome.

Smp_160700, a putative NSD1/NSD2 ortholog (Cabezas-Cruz et al., 2014), is another candidate involved in H3K36 trimethylation. It shows highest expression in cercariae, is 2-5 times higher expressed in different stages of the female juveniles and adults but is repressed in bO.

Smp_246410, a putative SETD5 ortholog (Padalino, 2019), may encode an enzyme involved in H3K36 trimethylation. This gene is highly expressed during larval stages and shows minimal sex-biased expression in the cell atlas, with only a slight increase in expression in bF compared to sF.

H3K9 di- and trimethylation are also conserved histone modifications. Based on homology searches, we confirmed that Smp_027300 (Cabezas-Cruz et al., 2014) and Smp_150850 (Padalino, 2019) as putative schistosome orthologs of SUV39H1/H2 and SETDB1/2, respectively. Smp_027300, is 2.6-fold higher expressed in ovaries compared to testes, and roughly 2 times higher expressed in bO than sO. Single-cell data indicate higher abundance in bF compared to sF. Smp_150850 has a similar expression pattern.

Due to the reversible nature of histone modifications, schistosome enzymes likely responsible for H3K36m3 or H3K9m3 demethylation could also be responsible for regulating this histone modification linked to gonadal maturation. JMJD2A is a Kme3-specific demethylase, reversing trimethylated H3K9 or H3K36 to dimethylated products. JMJD2A (also known as JHDM3A (Jumonji C (JmjC)-domain-containing histone demethylase 3A)) is a member of the JMJD2/KDM4 subfamily of demethylases and is evolutionarily conserved from *C. elegans* to Human. Using the beforementioned bioinformatics-led approach, we confirmed that Smp_132170, previously identified (Lobo-Silva et al., 2020), as the most likely enzyme responsible for H3K9me3/H3K36me3 demethylation. Smp_132170 has strongest expression in cercariae and is about 2 times more expressed in ovaries compared to testes. There is no difference between bO and sO or bT and sT, and mRNA is detected in many cell types in adult worm single-cell data.

In conclusion, there is evidence for pairing-dependent regulation of H3K9me in ovaries for Smp_210650 (a putative SETD2 catalysing trimethylation of H3K9 and H3K36), for Smp_027300 (a putative SUV39H1/H2), and for Smp_150850 (putative SETDB1/2). We hypothesized that these enzymes are involved in processes controlling oocyte differentiation, from the stem cell-like oogonia stage to the differentiated primary oocyte that has entered meiosis. Smp_246410 (putative SETD5 homologue) is a candidate for testis-specific H3K36 trimethylating activity.

In contrast, JMJD2/KDM4 homologue Smp_132170, coding for H3K9/38 demethylating activity is constitutively expressed. The details of these analyses are in supplementary data 9.

#### Blocking histone H3K9m3 and H3K36me3 de-methylation leads to pairing instability

Based on our results, we hypothesized that changing H3K9/K36 methylation status by inhibition of JMJD2/KDM4 homologue Smp_132170, a member of the demethylase family, would impede correct gonad functioning after pairing (Kooistra and Helin, 2012). To this end, we used a pharmacological inhibitor, IOX1, that was initially discovered by high-throughput screen against histone demethylase KDM4/JMDJ2 (https://www.thesgc.org/chemical-probes/IOX1). We incubated *S. mansoni* couples *in-vitro* with increasing concentrations of IOX1. Our data show a sharp increase in the separation of couples using ≥ 500 µM IOX1, which was not due to toxic effects as evaluated by vitality tests (supplementary data 10).

## Discussion

One of the central questions in developmental biology is how a developmental trajectory is maintained after the initial stimulus that had triggered a developmental switch has faded away. Epigenetic mechanisms are the usual suspects when developmental polymorphism is to be investigated. We define here “epigenetic” in the original terms of C. Waddington (Waddington, 1942) as markers that provide momentum to a developmental process leading to a defined alternative phenotype. One of the chromosomal epigenetic information carriers is post translational modification (PTM) of histones. We, therefore, decided to investigate histone modifications to better understand the molecular processes of *S. mansoni* gonadal changes upon pairing. To this end, we selected 8 histone modifications that had been linked to sexual differentiation and reproductive stem cell biology. Since the functional roles of PTMs are still elusive in *S. mansoni*, we first gained insight into the link between gene expression, chromatin structure and PTM by integrating RNA-Seq, ATAC-Seq and ChIP-Seq data. We then identified the most important PTM changes in isolated gonads of pairing inexperienced and experienced worms. This approach could also identify potential Achilles’ heels for new concepts to block gonad maturation and consequently egg production as strategies to fight schistosomiasis.

### A putative role of histone modifications in the three steps of transcription

About 60% of *S. mansoni* protein coding genes possess an enrichment of H3K4me3 around the TSS that increases relatively sharply upstream of the TSS and decreases progressively within the gene body. For these H3K4me3 positive genes we find a canonical pattern and, as in many other organisms, we see striking collinearity of histone PTM along protein coding genes with the different stages of transcription (Figure 7). H3K4me3 is found at the TSS, H3/H4 acetylation and, in single-sex testes H3K36me3, in the gene body, and H3/4 acetylation correlates with transcription strength. It is known that the C-terminal domain (CTD) of RNA polymerase II subunit Rpb1 undergoes dynamic phosphorylation, with different phosphorylation sites predominating at different stages of transcription (the “The RNA Polymerase II Carboxy-Terminal Domain (CTD) Code” (reviewed in (Eick and Geyer, 2013)). The enrichment of H3K4me3 at the TSS was a striking observation in all eukaryotes and several explanations for a link between CTD phosphorylation, PolII progression, and H3K4me3 were proposed. Binding of NuA3 histone H3 histone acetylase (HAT) and NuA4 H4 HAT complexes to H3K4me3 was found in yeast (reviewed in (Buratowski and Kim, 2010)), suggesting that H3K4m3 promotes acetylation in the downstream region increasing chromatin accessibility to PolII. A recent study in mouse cell cultures indicates that H3K4me3 is not required for transcription initiation, but regulates PolII pausing and potentially also has a role in elongation (Wang et al., 2023). For H3K36me3, results from yeast suggest a model of simultaneous binding of both the H3K36me “writer” (Set2) and “reader” (Eaf3/Rpd3S) to the CTD (Buratowski and Kim, 2010) providing a mechanistic link between histone code and CTD code. Interestingly, Pu et al (Pu et al., 2015) hypothesized that in *Drosophila melanogaster* and *Caenorhabditis elegans* little or no H3K36me3 marks in a gene might define a chromatin environment that provides greater flexibility in gene expression regulation over time. Therefore, it is possible that H3K36me3 may create a chromatin environment that allows these genes to stabilize transcription level. Increased acetylation of histones H3 and H4 is associated with enhanced transcriptional speed of PolII in genes. Likewise, higher H3K9 acetylation corresponds to a shift in the balance between distal and proximal pre-mRNA isoforms, indicating a localized acceleration of PolII (reviewed in (Muniz et al., 2021)). This goes in line with our observation that acetylation correlates with RNA-Seq abundance.

**Figure 7.**
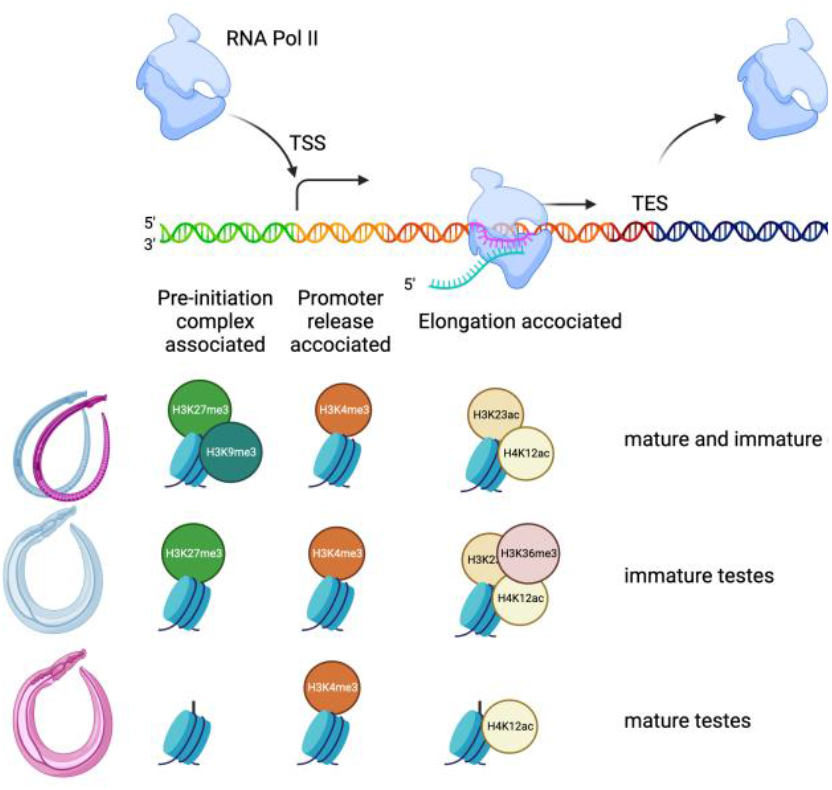
Putative relation of chromatin structure and transcription in *S. mansoni* gonads in H3K4me3-positive genes. In ovaries, H3K27me3 and H3K9me3 decorates the region upstream of genes that is pre-initiation complex associated. In male gonads, this is H3K27me3 in single-sex testes and an unknown PTM in mature testes. H3K4me3 sits in the transcription start site (TSS) and could be a signal for promotor release of the RNA PolII complex. Different histone acetylations are associated with the body of the gene and could be elongation associated. In sM testes, we also find H3K36me3 here. Made with BioRender.

A mark that differs from the canonical PTM distribution is H3K9me3 (and to a lesser degree H3K27me3) that shows a plateau upstream of the TSS with a sharp drop just upstream of the H3K4me3 peak. This was not only observed in the gonads but also in the other live cycle stages of *S. mansoni* (Roquis et al., 2015). We verified that this peculiarity is not due to a mistake in our analysis pipeline by re-analyzing published human ChIP-Seq data (supplementary file 11). Clearly, *S. mansoni* and its human host differ in the use of H3K9me3 gene decoration. In many organisms, H3K9me3 can recruit Heterochromatin Protein 1 (HP1), and it was suggested that HP1 locally increase Pol II pausing on alternatively spliced exons enriched in H3K9me3 marks, and that H3K27me3 decrease PolII speed (reviewed in (Muniz et al., 2021)). We showed earlier (da Trindade et al., 2025) that in cercariae, *Sm*HP1 is enriched in transcription end sites but has also a smaller peak at TSS. It is conceivable that *S. mansoni* uses H3K9me3/HP1 to slow down PolII to enable the formation of the pre-initiation complex. Further work is needed to address the question. Nevertheless, for annotation purpose or the determination of genomic safe harbours, it is noteworthy that the H3K9me3 peak followed by H3K4me3 peak in 5’ to 3’ direction is indicative of direction and strength of transcription.

For H3K4me3-negative genes, there is no clear co-linearity with the stages of transcription. It might be that in these genes, *Schistosoma*-specific histone PTMs are associated with PolII, a hypothesis that warrants further investigation by *de-novo* PTM discovery approaches.

### The different roles of H3K4me3 positive and negative protein coding genes

About 40% of *Schistosoma* genes possess no H3K4me3 in their TSS while being transcribed. A similar situation has been observed in other species such as budding yeast (Murray et al., 2019) and *Drosophila* (Hödl and Basler, 2012) raising the question whether H3K4me3 is a transcriptional activator? Our findings indicate that *S. mansoni* genes possess H3K4me3 in their TSS, and H3K4me3 occupancy is indeed the best predictor for expression strength. In other words, as more H3K4me3 is present, more mRNA will be produced (or *vice versa*). However, this does not mean that genes without H3K4me3 in their TSS are not transcribed, even though their level of transcription is, on average, lower. We show that not only H3K4me3 occupancy, but also the histone modification profiles along genes, are different between H3K4me3-negative and positive genes.

This poses the question why schistosomes may have evolved two sets of chromatin colours? It could be a way to tag housekeeping and inducible genes, but here we show that H3K4me3 does not differentiate inducible *vs* constitutively expressed genes in schistosome gonads. Instead, GO enrichment analysis suggests that H3K4me3 is used by the parasite to control genes that code for functions being evolutionary ancient and located in the interior parts of the cells. In contrast, functions that are necessary to communicate with the outside, including the host environment (or the partner during male-female interaction), are painted with other, so far unknown histone modifications. It is noteworthy that among the H3K4me3-negative genes there are (i) all micro-exon genes (MEGs) that are candidates for interaction with hosts, but also (ii) Smp_158480 the *Schistosoma mansoni non*-*ribosomal-peptide synthetase* (*Sm-nrps)* (Chen et al., 2022), Smp_135230 the *L-tyrosine decarboxylase* (Sm*tdc-1*), and Smp_171580 the *DOPA decarboxylase* (Sm*ddc-1*) that are known to influence female sexual maturation in a pairing-dependent manner (Li et al., 2023). Among the H3K4me3-negative genes of potential importance for sexual biology is also the homologue of doublesex/male-abnormal-3 domain transcription factor *Sm_dmd-1* Smp_143190 that is highly expressed in the testis (https://v7test.schisto.xyz/smansoni-v7/smp_143190/). This gene was proposed to be important for expression of male-specific functions by homology with a free living a simultaneous hermaphrodite, the planarian *Schmidtea mediterranea* (Chong et al., 2013).

We hypothesise that this functional compartmentalisation has evolved due to a fundamental constraint of parasitism: host tissue constitutes the parasite’s immediate environment, and histones together with their modifying enzymes are evolutionarily ancient and highly conserved across clades. As a result, ancestral schistosomes likely possessed histone-modifying enzymes (epigenetic “writers”) and epigenetic “readers” similar to those of their hosts, allowing hosts to “read” and potentially respond to the parasite’s histone code and/or *vice versa*. A plausible adaptive response would be the evolution of an alternative code by the parasite. We therefore propose that, at genes lacking H3K4me3, contemporary schistosomes deploy a distinct, parasite-specific epigenetic code that is less accessible to host decoding. A schematic of this model is shown in Figure 8. Consistent with this hypothesis, we observe that H3K9me3 is used differently in schistosomes than in their human hosts, and our prior finding that lncRNA genes exhibit non-canonical histone post-translational modification profiles (Augusto et al., 2025) also fits this conceptual framework.

**Figure 8.**
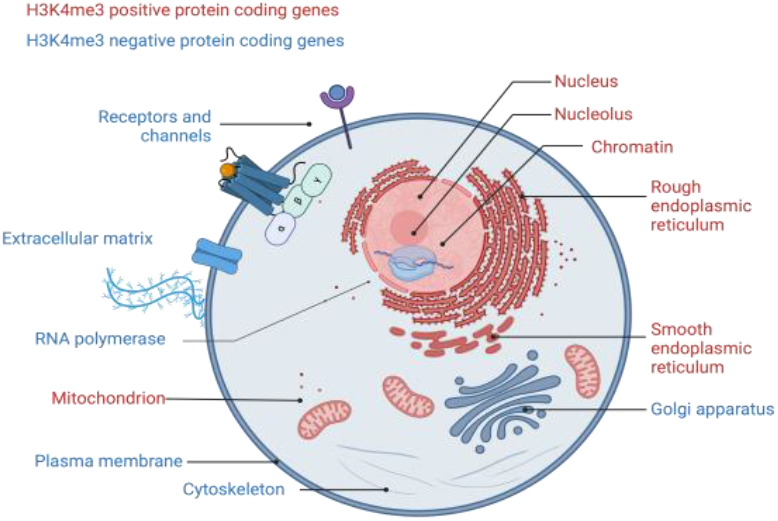
Schematic representation of GO enrichment analyses results for H3K4me3 positive genes (red) and H3K4me3 negative genes (blue). Genes that are enriched in functions and cellular locations that serve to interact with the host environment are lacking H3K4me3 and there might be schistosome-specific histone modifications decorating them. Figure created with BioRender.

### Mayor changes in histone acetylation and H3K9 and H3K36 trimethylation occur in gonads after pairing

Changes in histone modifications after pairing are massive, covering up to 50% of the genome for some modifications and up to tens to hundreds of thousand regions, depending on how differences are defined. In ovaries, these changes occur predominantly in H3K23ac, H3K9ac, and H3K36me3. Similar to the ovaries, major changes are observed in H3K23ac, H3K9ac, H4K12ac in the testes, but instead of H3K36me3, it is H3K9me3 that strongly changes its distribution during the transition from pairing inexperienced to experienced status. To put this into perspective, we can compare these results with earlier published comparison between cercariae and adults (Roquis et al., 2018). Here, we had investigated by ChIP-Seq only H3K4me3, H3K27me3, and H4K20me1 and identified 1,195 differences in histone PTM occupancy with 552 contributed by H3K4me3 alone. This is different in gonads where H3K4me3 remains stable (sT vs bT: 99, sO vs bO: 107). In contrast, we see several thousand differences in H3K27me3 and H4K20me1 (table 3), while it was only a couple of hundred between cercariae and adults (Roquis et al., 2018). These data are not directly comparable because in the earlier study whole organisms were used and not isolated organs. Therefore, the higher variance observed in the 2018 study shows that histone PTM changes during gonad maturation is tremendous.

Given the important changes in histone PTMs after pairing, we wondered what blocking the underlying enzymatic activity would produce as effect.

We had had earlier shown that histone methyltransferase inhibitors block miracidium to primary sprorocyst transformation (Roquis et al., 2018), and that histone deacetylase inhibitors trichostatin A (TSA) reversibly blocks metamorphosis of miracidia into sporocyst (Azzi et al., 2009). We choose treatment with IOX1 that targets the de-methylation activity of KDM4/JMDJ2 and, therefore, was expected to impact both H3K36 and H3K9 methylation. Treatment leads to separation of mated worms without observable toxic effects indicating that H3K36/9 methylation status is important for maintaining the couple together. Since IOX1 disrupts pairing after it had been correctly established, the activity of the KDM4/JMDJ2 homologue must be continuous to ensure the maintenance of the mating state—mere initiation (“love”) is not sufficient for persistence. This interpretation aligns well with single-cell transcriptomic data, which show that the KDM4/JMDJ2 homologue Smp_132170 is broadly transcribed across cell types. We did not find data on knockdowns in the literature. Besides being of academic interest, our results identified H3K9 and H3K36 (de)methylation also as a potential target for drug development since divorce would impede egg production as one of the pathological mechanisms and also interrupt the life cycle of the parasite. A major challenge in drug development targeting histone-modifying enzymes in schistosomes is their high homology to human counterparts, raising concerns about off-target effects. But subtle structural differences might still be enough to produce selective drugs (Pizarro et al., 2013). Interestingly, our results also show that the full repertoire of histone modifications in schistosomes remains largely unexplored, and it is likely that many remain unidentified. These limitations underscore the need for a *de novo* approach aimed at discovering schistosome-specific histone modifications, which could reveal novel, selective targets for therapeutic intervention.

## Materials and Methods

### Ethics statement

Housing, feeding and animal care at Justus Liebig University Giessen, Germany, followed the national ethical standards setting the conditions for approval, planning and operation of establishments, breeders and suppliers of animals used for scientific purposes and controls. The animal experiments were approved by the Regional Council Giessen (V54-19 c 20/15 c GI 18/10).

### Biological material

Gonad isolation was performed by a detergent and enzyme-based isolation procedure (Hahnel et al., 2013). In short, worm couples were separated immediately after perfusion, and about 50 (males and bisex females) or 100 (single-sex females) individuals were transferred into 2ml Eppendorf tubes, respectively, and washed with 2 ml M199 (non-supplemented) medium. Tegument removal was achieved by incubating worms for 2× 5min (females) or 3× 5min (males) in 400μl stripping buffer, respectively (0.1% of each: Brij 35 (Roth), Nonidet P40 (NP-40, Fluka), Tween 80 (Sigma), and Triton X-405 (Sigma) in DEPC-PBS, pH 7.2–7.4; 0.2 μm filtered before use) at 1,200 rpm and 37 °C on a thermal shaker (Biosan TS-100, Latvian Republic). Afterwards, the worms were washed with 2ml M199 (non-supplemented, 3x), the medium removed, and enzymatic digestion started by incubating each worm sample with 300–500 μl elastase (Sigma E0258; 5U/ml in non-supplemented M199) for 20– 40min at 650 rpm and 37 °C. The process of tissue digestion was checked microscopically (Leica). The tube contents were transferred to 2.5 cm Petri dishes filled with M199 (2 ml each). Intact gonads were collected by pipetting with 10 μl tips, and transferred 2–3 times to new dishes for purification. Organs collected this way were transferred into RNase-/DNAse-free vessels on ice, centrifuged at 6,000g for 2min and frozen at −80°C after removing the supernatant. In parallel, 5–10 organs were stained 10μl Trypan Blue (0.4%; Sigma) for vitality check with similar results as described before (Hahnel et al., 2013).

### Chromatin immunoprecipitation and sequencing (ChIP-Seq)

We performed native chromatin immunoprecipitation followed by sequencing (ChIP-Seq) as described for *S. mansoni* in (de Carvalho Augusto et al., 2020). For each histone mark, we had two biological replicates for testis and ovaries from immature (sM and sF) and mature (bM and bF) adult worms. Immunoprecipitation was performed using antibodies against H3K4me3, H3K27me3, H3K9me3, H3K9ac, H3K23ac, H4K20me1, H4K12ac, H3K36me3 and details for each antibody (supplier, lot number, amount used) are in (Augusto et al., 2025).

For each sample, we used a control without antibody to assess nonspecific background (bound fraction) and input (unbound fraction). All antibodies were carefully tested for specificity as described (Cosseau and Grunau, 2011). Immunoprecipitated DNA were quantified on a high sensitivity bioanalyser, before being cleaned with Agencourt AMPure XP beads. End repair, A-tailing and adapter ligation were performed using the NEB library prep kit, with Agencourt AMPure XP bead cleaning steps between each stage. The amount of template for PCR and the number of PCR cycles required were assessed from a high sensitivity bioanalyser trace post-ligation. Libraries were amplified with 14 cycles. After cleaning with Agencourt AMPure beads, libraries were quantified using a KAPA SYBR FAST ABI Prism qPCR Kit with Illumina GA Primer Premix (10x) and 7 x Illumina GA DNA Standards (Kit code: KK4834) on an ABI StepOnePlus qPCR machine. Libraries were diluted into an equimolar pool and run on a HiSeq 2500, generating 75 base pair, paired end reads on an Illumina HiSeq 2500 at Wellcome Trust Sanger Institute (UK).

### Quality control, alignment, and peak calling

All data processing was performed on a local GALAXY instance (http://bioinfo.univ-perp.fr) (Giardine et al., 2005). Read quality was verified using the FastQC toolbox (https://www.bioinformatics.babraham.ac.uk/projects/fastqc/). All samples had a minimum average read quality score of 30 over 95% of their length, and no further cleaning steps were performed.

Sequences were aligned to the *S. mansoni* reference genome v7 (Protasio et al., 2012) with Bowtie v2.1 (Langmead and Salzberg, 2012) using parameters–end-to-end,–sensitive,– gbar 4. BAM files generated by Bowtie2 were coordinate sorted and then filtered for unique matches using the XS tag with samtools v1.3.1 (Li et al., 2009) (samtools view -Sh -q quality value 40–42—F 0×0004 –| grep -v XS:i). PCR duplicates were also removed using samtools (samtools rmdup). Although not mandatory, we found that performing random sampling to use the same amount of uniquely mapped reads for each sample and each histone mark improved sensitivity and specificity when looking for chromatin structure differences. Peak calling using PEAKRANGER v1.16 (Feng et al., 2011) (parameters: P-value ≤ 0.0001, False discovery rate (FDR) ≤ 0.01, read extension length 200, smoothing bandwidth 99 and Delta 1) was used to visualize histone mark distributions. Input samples were used as a negative control (-c) for normalization. Wiggle files generated in the process were visualized with IGV (Thorvaldsdóttir et al., 2013).

### Chromatin landscape of unique sequences

We started our analyses by characterizing the genome-wide chromatin landscape in for testis and ovaries from immature (sM and sF) and mature (bM and bF) of *S. mansoni*. We used ChromstaR (v1.2.0) for peak detection and comparative analysis, using epialleles that ChromstaR detected in all replicates of a given developmental stage. ChromstaR is an R package that uses Hidden Markov Models (HMM) to perform computational inference of discrete combinatorial chromatin state dynamics over the whole genome (Taudt et al., 2016). It can perform uni- or multivariate analyses using several replicates and identifying common and different peaks between conditions. ChromstaR was processed in two steps: (1) we fitted a univariate Hidden Markov Model over each ChIP-seq sample individually (*i.e*. each replicate for each condition) and (2) we performed a multivariate HMM over the combined ChIP-seq samples. BAM files from each condition were processed under the combinatorial mode, with a false discovery rate (FDR) cutoff of 0.05 and bin size of 150 and the input for background correction. Transcriptional start sites (TSS) of genes were defined as 3 bp (one base pair upstream and downstream of the +1 transcription site) based on the genome annotation file v.7 downloadable from ftp://ftp.sanger.ac.uk/pub/pathogens/Schistosoma/mansoni/genome/GFF/. A simplified BED file with TSS and transcription end sites (TES) was generated for contigs assembled at chromosome level. Chromosome names were changed to Chr1—ChrW. Knowing that histone modifications have crucial roles in the regulation of gene expression, we tested the genome-wide coverage (*i.e*. the percentage of the genome that is covered by the chromatin state) of each histone mark over the genome and at TSS. In the following, chromatin profiles were generated for fragments over several base pairs (bp) in length.

### Chromatin landscape of repetitive sequences

ChIP-Seq analyses were performed as described in (Roquis et al., 2018). FASTA file of consensus repeats also in supplementary file 12.

### Association between RNA-Seq and ChIP-Seq data

We built a random forest (R package v4.6-14) classifier to explore the association between RNA-seq gene expression and levels of eight histone modifications (counts of histone marks, compiled RNA-Seq and ChIP-Seq raw data in supplementary file 1). Firstly, to obtain gene expression data, we mapped the raw RNA-seq data of *S. mansoni* gonads (Lu et al 2016) to the updated Sm_V7 genome (WormBase Parasite 14) using STAR v2.4.2a and summarized read counts using featureCounts v1.4.5-p1, a pipeline described before (Lu et al., 2017), https://v7test.schisto.xyz/dataused/). The raw counts were processed using edgeR 3.28.1 to obtain normalized counts and RPKM values, which were used to quantify gene expression in this study. Differentially expressed genes between mature and immature gonads were identified using DESeq2 1.26 with the option cooksCutoff=TRUE and threshold set at padj < 0.05 and at least 1.5-fold change. To obtain the counts of individual histone marks (peaks) in various genomic regions, we first defined target regions for each histone mark as followings: 1) promoter region (2kb upstream TSS and 200bp downstream TSS) for H3K9me3; 2) near TSS (200bp upstream TSS to the middle of the gene) for H3K4me3, H4K20me1, and H3K27me3; 3) gene body region for H3K9ac, H3K23ac, and H4K12ac; 4) towards TES (middle of the gene to 200bp downstream TES) for H3K36me3. The gene coordinates were also obtained based on Sm_V7 annotation and histone marks were summarized using the command bedtools intersect with PEAKRANGER output bed files at binsize 150 and stepsize 75. Based on these steps we generated the supplementary file 1 for further analysis. For random forest modelling, we first filtered out genes without any histone mark but still expressed at rpkm>1 as potential outliers. The model was built for each sample using histone marks as features and log2-transformed rpkm values as observations, with 80% genes for training and 20% for testing, using 2000 trees and setting importance=TRUE. We also run the modelling for genes with and without H3K4me3 marks separately. In the testing set, Pearson’s correlation between predicted and true expression, as well as the Root Mean Seqare Error (RMSE) values were calculated to evaluate the model performance.

To evaluate the association between gene expression and chromatin colours, we used bedtools intersect to summarize the counts of different histone combinations (colours) in the region of 2kb upstream TSS and 200bp downstream TES, based on binsize 500 and stepsize 250 outputed peak bed files. Similarly, genes without any detected colour but with rpkm>1 were regarded as outliers and removed. A random forest classifier was built with 2000 trees using 80% genes as training and the rest 20% as testing.

### Gene ontology enrichment analysis

GO file that associates Smp_ numbers with Gene Ontology IDs was downloaded from https://parasite.wormbase.org/index.html. We then used Rank-based Gene Ontology Analysis with Adaptive Clustering (RbGOA) (https://github.com/z0on/GO_MWU) which was initially developed to measure whether each GO category is significantly enriched by either up or downregulated genes. Here, we used H3K4me3 peak strength instead of expression data. GO terms that best represent independent groups of significant GO terms identified by RbGOA were used for graphical representation.

### Use of metagene profiles for clustering genes into chromatin types

Metagene profiles were exported from ChromstaR as individual TSV files. Under R, a loop was used to iterate over each file name in the file_names vector, load the corresponding distance matrix from the file using read.csv, convert it to a matrix using as.matrix, and append it to the dist_matrices list using append. After loading all the distance matrices, the code used *Reduce* along with the + operator to combine the distance matrices element-wise into the combined_dist_matrix. Hierarchical clustering was performed using the *hclust()* function, which takes the combined distance matrix as input. The *as.dist()* function is used to convert the matrix into a distance object required by *hclust()*. After performing hierarchical clustering, the resulting dendrogram was plotted using the *plot()* function. The R scripts are available in the “Scripts” section of the supplementary files.

### Clustering of metagene profiles into similar shapes (“types”)

Metagene profiles over all 256 combinations were exported from ChromstaR as individual TSV files for each condition and converted into TSV files with one column per chromatin colour and the header column relative positions around genes (+/-2000 kb of TSS end TES) with a custom script (*Convert chromstar output to tab by marks.py*). The resulting file was edited to remove the first column, remove duplicated rows “0” and “2000” and to replace “-Inf” by “0”. We then transposed the data matrix and performed hierarchical clustering on the data with the *hclust* R function. A dendrogram was plotted to guide the decision on the number of metagene types and a table of the colour names and their cluster numbers exported. Colours were sorted into clusters, we kept only the 54 biological relevant chromatin colours and totalled the metagene values for each of the chromatin colours belonging to the same metagene type for visual representation as a metagene profile. The R scripts are available as compressed archive in supplementary file 12.

## End Matter

### Author Contributions and Notes

CG, RCA, MB, CGG and KFH designed research, CG, RCA, ZL, AC, MM, TQ, CC, EA, JFA, AR, NH, GP performed research. ZL and AC wrote software. ZL, AC, CC, CG, RCA, EA, AR analyzed data, and all authors wrote the paper.

The authors declare no conflict of interest.

This article contains supporting information online: raw data are available under the NCBI SRA BioProject accession PRJNA1173904. supplementary files are at Zenodo https://doi.org/10.5281/zenodo.16745298.

## Acknowledgments

This study received financial support from the Wellcome Trust strategic award 107475/Z/15/Z (FUGI). With the support of LabEx CeMEB, an ANR « Investissements d’avenir » program (ANR-10-LABX-04-01) and the Environmental Epigenomics Core Service at IHPE. This study is set within the framework of the « Laboratoire d’Excellence (LabEx) » TULIP (ANR-10-LABX-41). Funding was provided by the French National Research Agency (ANR) under grant agreement ANR-22-CPJ1-0056-01, within the framework of the project “Tropical diseases of today, European diseases of tomorrow: a systems biology approach to understand, predict and control their emergence”.

